# A survey on antimicrobial resistance genes of frequently used bacteria in kefir and yoghurt

**DOI:** 10.1101/2022.01.12.476061

**Authors:** Maura Fiona Judge, Adrienn Gréta Tóth, Sára Ágnes Nagy, Márton Papp, Norbert Solymosi

## Abstract

Antimicrobial resistance (AMR) is one of the foremost threats facing the treatment of infectious diseases worldwide. Recent studies have highlighted the potential for antimicrobial resistance genes (ARGs) in fermented foods to contribute to AMR via horizontal gene transfer (HGT). The focus of our study was investigating the ARG content (resistome) and mobility potential of the ARGs (mobilome) of bacterial strains commonly used in probiotic products, namely yoghurt and kefir. We performed metagenomic analyses on freely available data sets (n=584) originating from various kefir and yoghurt strains using next generation sequencing (NGS) in order to gain an insight into the ARG diversity, frequency and mobility. Our study shows that kefir and yoghurt products carry diverse and significant amounts of ARGs and that these genes may often be associated with iMGEs or plasmids, conferring mobility. Certain bacteria species such as *Bifidobacterium animalis* and *Streptococcus thermophilus* were found to have higher ARG content. Overall, our results support the hypothesis that ARGs are present in fermented foods, namely yoghurt and kefir, and have the potential to contribute to AMR.

## Introduction

Antimicrobial resistance (AMR) is one of the foremost threats facing the treatment of infectious diseases worldwide, both in human and animal medicine. Dealing with the COVID-19 pandemic of late highlights the significance of this threat. Antimicrobials are used to treat human and animal diseases, and since antimicrobial use has been increasing, so too has AMR causing these antimicrobials to become less effective. It is currently estimated that 700,000 people are dying per annum from AMR related issues, with projections forecasting this number to rise to 10 million by 2050^1^. Identifying potential sources of AMR are thus of utmost importance. AMR can be acquired by bacteria via gene mutations or horizontal gene transfer (HGT)^2^. HGT occurs primarily by transformation, conjugation or transduction and involves small packets of DNA being transferred between bacteria. The transfer of antimicrobial resistance genes (ARGs) is enhanced by being linked with mobile genetic elements (MGEs), particularly plasmids^3, 4^. In this study, as a continuation of our work^5, 6^, we explored the possible development of AMR due to HGT during the fermentation of food produce derived from animal sources, namely kefir and yoghurt products. The multiplication of bacteria during the fermentation process is widely known and understood. However, a less studied side to this is that if their genomes harbour ARGs, then their amount is increasing too, which could potentially be aiding the development of AMR.

Probiotics are encouraged to restore natural microbiomes and a healthy gut^7^. However, there is some question over their effectiveness^8, 9^. Instead of in-situ samples, studies mostly rely on stool samples, which have been shown to often be inaccurate representations of the gut microbiome^10, 11^. In addition, there has been some concern over the effects of the bacteria that probiotics harbour^12^. Recent studies indicate that the genomes of the bacterial composition of fermented foods contain ARGs^13–17^. The gut microbiome may thus act as a resistome^18^. This would give ARGs, even from non-pathogenic bacteria, the opportunity to spread via HGT to pathogenic bacteria that they become physically close to, creating ‘superbugs’. In this study, we looked to further explore whether probiotics contribute to this resistome and thus possibly to the development of AMR worldwide. I collated data from other studies to identify the most frequently found bacterial species in kefir and yoghurt products. Using this data, we explored the ARG content of eight bacterial species by performing metagenomic analyses on freely available data sets originating from kefir and yoghurt strains. Using next generation sequencing (NGS) we were able to gain an insight into the ARG diversity, frequency and mobility of those data sets.

## Materials and Methods

### Data

Published studies^19–24^ were collated to determine the most frequently identified bacterial strains found in kefir and yoghurt products (Supplementary table). Since we have found high numbers of species in the literature, we selected the following most common ones for further analyses: *Bifidobacterium animalis, Lacticaseibacillus casei, Lacticaseibacillus paracasei, Lactiplan-tibacillus argentoratensis, L. plantarum, Lactobacillus helveticus, Levilactobacillus brevis* and *Streptococcus thermophilus*. All the available suitable samples were collected for these species from the National Center for Biotechnology Information (NCBI) Sequence Read Archive (SRA) repository. During the search I checked if the reads were suitable using predetermined parameters. The finding were filtered for the source by DNA and the platform by Illumina. Only those results with WGS or WGA strategies, genomic sources, random or PCR selections and paired layouts and at least a million clusters were selected. The list of analysed NCBI SRA run identifiers can be found in the supplementary materials.

### Bioinformatic analysis

Quality based filtering and trimming of the raw short reads was performed with TrimGalore (v.0.6.6, https://github.com/FelixKrueger/TrimGalore), setting 20 as a quality threshold. Only reads longer than 50 bp were retained. The preprocessed reads were assembled to contigs with MEGAHIT (v1.2.9)^25^ using default settings. From the contigs all possible open reading frames (ORFs) were gathered with Prodigal (v2.6.3)^26^. The protein translated ORFs were aligned to the ARG sequences of the Comprehensive Antibiotic Resistance Database (CARD, v.3.1.3)^27^, ^28^ by Resistance Gene Identifier (RGI, v5.2.0) with Diamond^29^. The ORFs classified as perfect or strict were further filtered with 90% identity and 60% coverage. All nudged hits were excluded. Integrative mobile genetic element (iMGE) content of contigs harbouring ARG was analyzed with MobileElementFinder (v1.0.3) and its database (v1.0.2).^4^ Following the distance concept of Johansson et al.^4^ for each bacterial species, only those with a distance threshold defined within iMGEs and ARGs were considered associated. The plasmid origin probability of the contigs was estimated by PlasFlow (v.1.1)^30^ The phage content of the assembled contigs was prediced by VirSorter2 (v2.2.1)^31^. The findings were filtered for dsDNAphages and ssDNAs. All data management procedures, analyses and plottings were performed in R environment (v4.1.0).^32^

## Results

The analysis of the short read datasets (n=584) from the metagenomic samples of *Bifidobacterium animalis, Lacticaseibacillus casei, L. paracasei, Lactiplantibacillus argentoratensis, L. plantarum, Lactobacillus helveticus, Levilactobacillus brevis* and *Streptococcus thermophilus* are summarised in the two following sections; Resistome and Mobilome. Following the presentation of the identified ARGs (resistome), the mobility potential of the ARGs (mobilome) is summarized based on the identification of iMGEs in the sequence context of the ARGs and contigs harbouring the ARGs being identified as plasmid originated. We did not find any phage-associated ARGs.

### Resistome

The resistome results are summarized in Table 1 and Figures 1 and 2. *Bifidobacterium animalis* had the lowest diversity of ARGs with two distinct ARGs identified and *Streptococcus thermophilus* had the highest with twenty-seven.

**Table 1.**
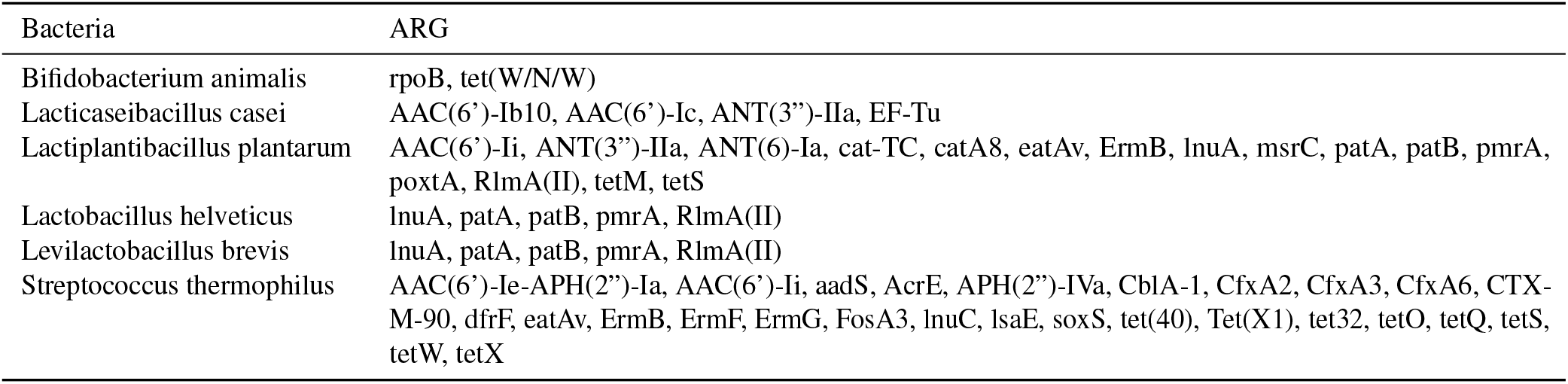
Identified ARGs by species. The gene names that are too long have been abbreviated (cat-TC: *Lactobacillus reuteri* cat-TC; EF-Tu: *Escherichia coli* EF-Tu mutants conferring resistance to Pulvomycin; rpoB: *Bifidobacterium adolescentis* rpoB mutants conferring resistance, to rifampicin; soxS: *Escherichia* coli soxS with mutation conferring antibiotic resistance)

| Bacteria | ARG |
| --- | --- |
| *Bifidobacterium animalis* | rpoB, tet(W/N/W) |
| *Lactocaseibacillus casei* | AAC(6')-Ib10, AAC(6')-Ic, ANT(3'')-IIa, EF-Tu |
| *Lactiplantibacillus plantarum* | AAC(6')-Ii, ANT(3'')-IIa, ANT(6)-Ia, cat-TC, catA8, eatAv, ErmB, lnuA, msrC, patA, patB, pmrA, poxtA, RlmA(II), tetM, tetS |
| *Lactobacillus helveticus* | lnuA, patA, patB, pmrA, RlmA(II) |
| *Levilactobacillus brevis* | lnuA, patA, patB, pmrA, RlmA(II) |
| *Streptococcus thermophilus* | AAC(6')-Ie-APH(2'')-Ia, AAC(6')-Ii, aadS, AcrE, APH(2'')-IVa, CblA-1, CfxA2, CfxA3, CfxA6, CTX-M-90, dfrF, eatAv, ErmB, ErmF, ErmG, FosA3, lnuC, lsaE, soxS, tet(40), Tet(X1), tet32, tetO, tetQ, tetS, tetW, tetX |

**Figure 1.**
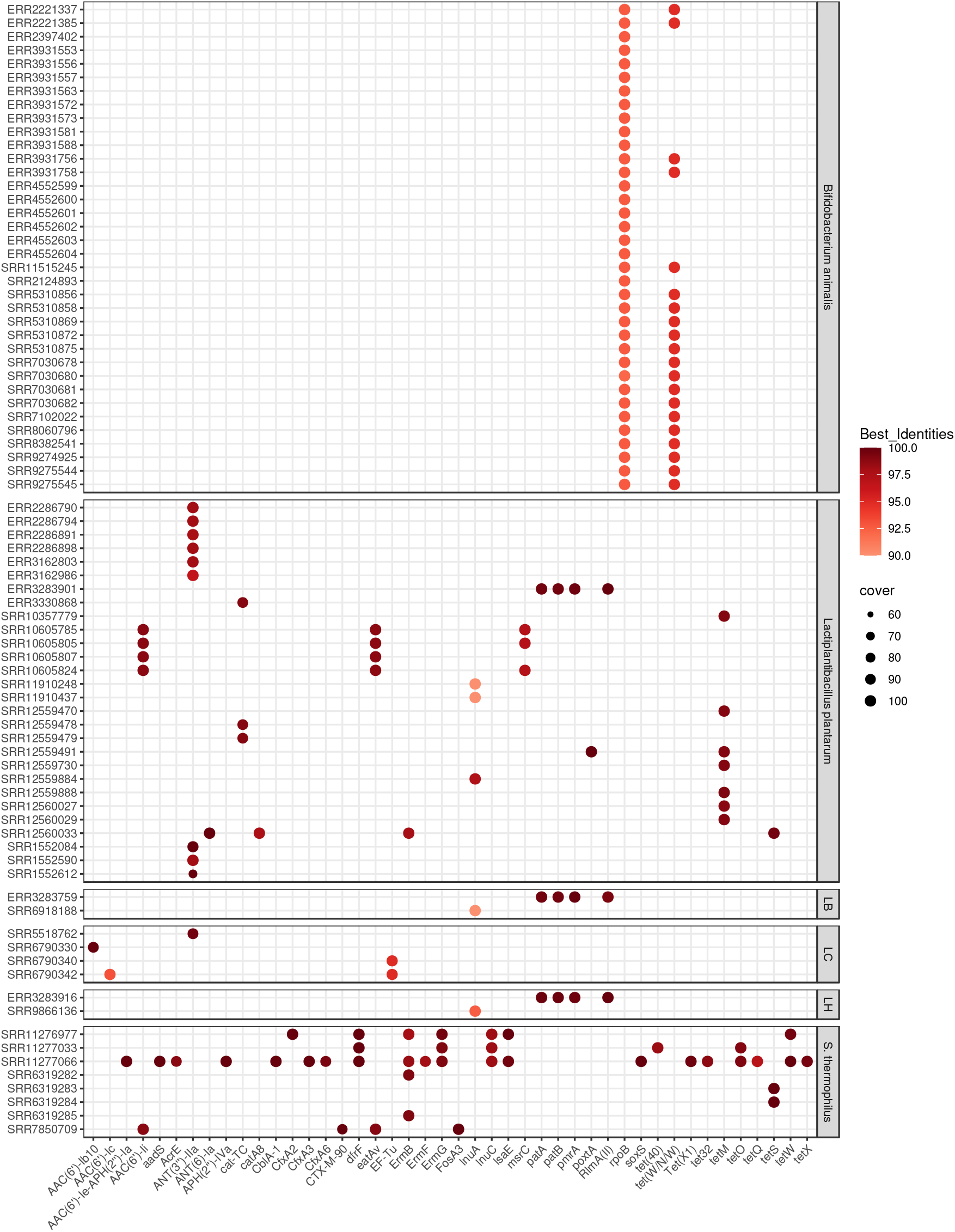
Identified ARGs by sample. For each sample-ARG combination, only the best finding is plotted. The size of the dot corresponds to the coverage and the colour to the sequence identity of hits on reference genes. Abbreviated species names: LB: *Levilactobacillus brevis*; LC: *Lacticaseibacillus casei*; LH: *Lactobacillus helveticus*. Gene names which are too long have been abbreviated (cat-TC: *Lactobacillus reuteri* cat-TC; EF-Tu: *Escherichia coli* EF-Tu mutants conferring resistance to Pulvomycin; rpoB: *Bifidobacterium adolescentis* rpoB mutants conferring resistance to rifampicin; soxS: *Escherichia coli* soxS with mutation conferring antibiotic resistance).

**Figure 2.**
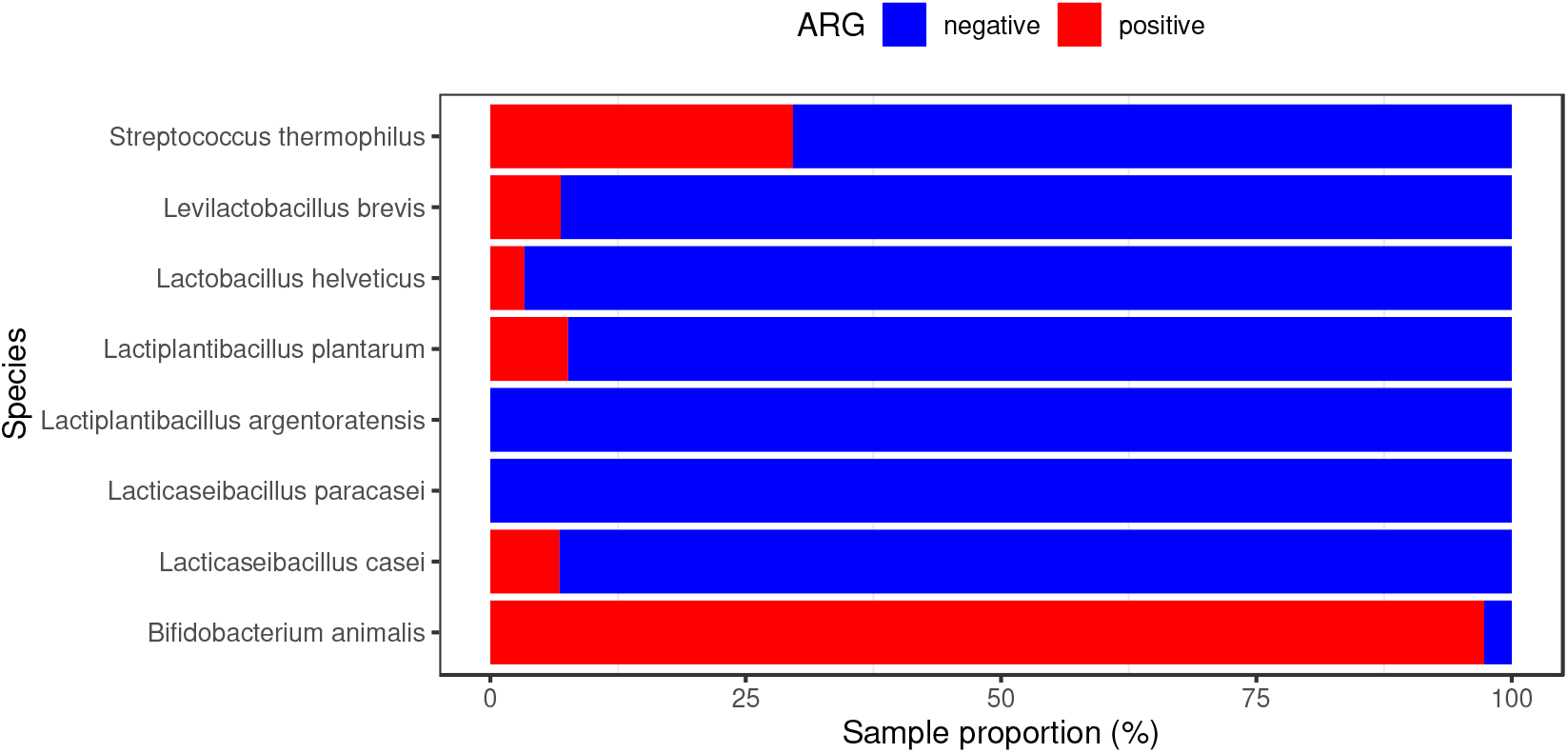
Antimicrobial resistance gene (ARG) positive sample frequencies. The proportion of ARG positive samples by species.

Figure 1 illustrates all identified ARGs by species and sample. The size of dot indicates coverage (the proportion of the reference ARG sequence covered by the ORF). The colour indicates the percentage sequence identity to the reference ARG. The most common ARGs were rpoB mutants conferring resistance to rifampicin and tet(W/N/W) which were detected in 36 and 20 samples respectively, all in Bifidobacterium animalis. The number of distinct ARGs found within a given sample ranged from 1 to 20 with the highest number of ARGs found in sample SRR11277066 obtained from Streptococcus thermophilus.

Figure 2 represents the proportion of positive and negative ARG samples of each of the eight bacterial species. These results show that all species contained ARGs except Lactiplantibacillus argentoratensis and Lacticaseibacillus paracasei. Levilactobacillus brevis, Lactobacillus helveticus, Lactiplantibacillus plantarum and Lacticaseibacillus casei all had a low proportion of ARG positive samples. This is in contrast with Bifidobacterium animalis which had a higher proportion of ARG positive samples.

The ARGs belonging to the genome of *Bifidobacterium animalis* may play a role in the appearance of resistance against rifamycin and tetracycline; *Lacticaseibacillus casei*: aminoglycoside, elfamycin; *Lactiplantibacillus plantarum*: aminoglycoside, fluoroquinolone, lincosamide, macrolide, oxazolidinone, phenicol, pleuromutilin, streptogramin, tetracycline; *Lactobacillus helveticus*: fluoroquinolone, lincosamide, macrolide; Levilactobacillus brevis: fluoroquinolone, lincosamide, macrolide; *Streptococcus thermophilus*: aminoglycoside, carbapenem, cephalosporin, cephamycin, diaminopyrimidine, fluoroquinolone, fosfomycin, glycylcycline, lincosamide, macrolide, monobactam, oxazolidinone, penam, penem, phenicol, pleuromutilin, rifamycin, streptogramin, tetracycline, triclosan.

The proportions of resistance mechanisms were calculated based on the ARG diversity. The dominant mechanisms of identified ARGs were antibiotic target protection (30.72%), antibiotic inactivation (26.8%), antibiotic target alteration with antibiotic target replacement (23.53%), antibiotic target alteration (9.15%), antibiotic efflux (7.19%), antibiotic target replacement (1.96%) and antibiotic target alteration with antibiotic efflux and reduced permeability to antibiotics (0.65%).

### Mobilome

The frequency of iMGEs and plasmids associated with the ARGs is summarized by bacteria of origin in Figure 3. This figure represents the mobility of the ARGs identified. The size of the dot indicates the number of occurrences of the given mobile ARG.

**Figure 3.**
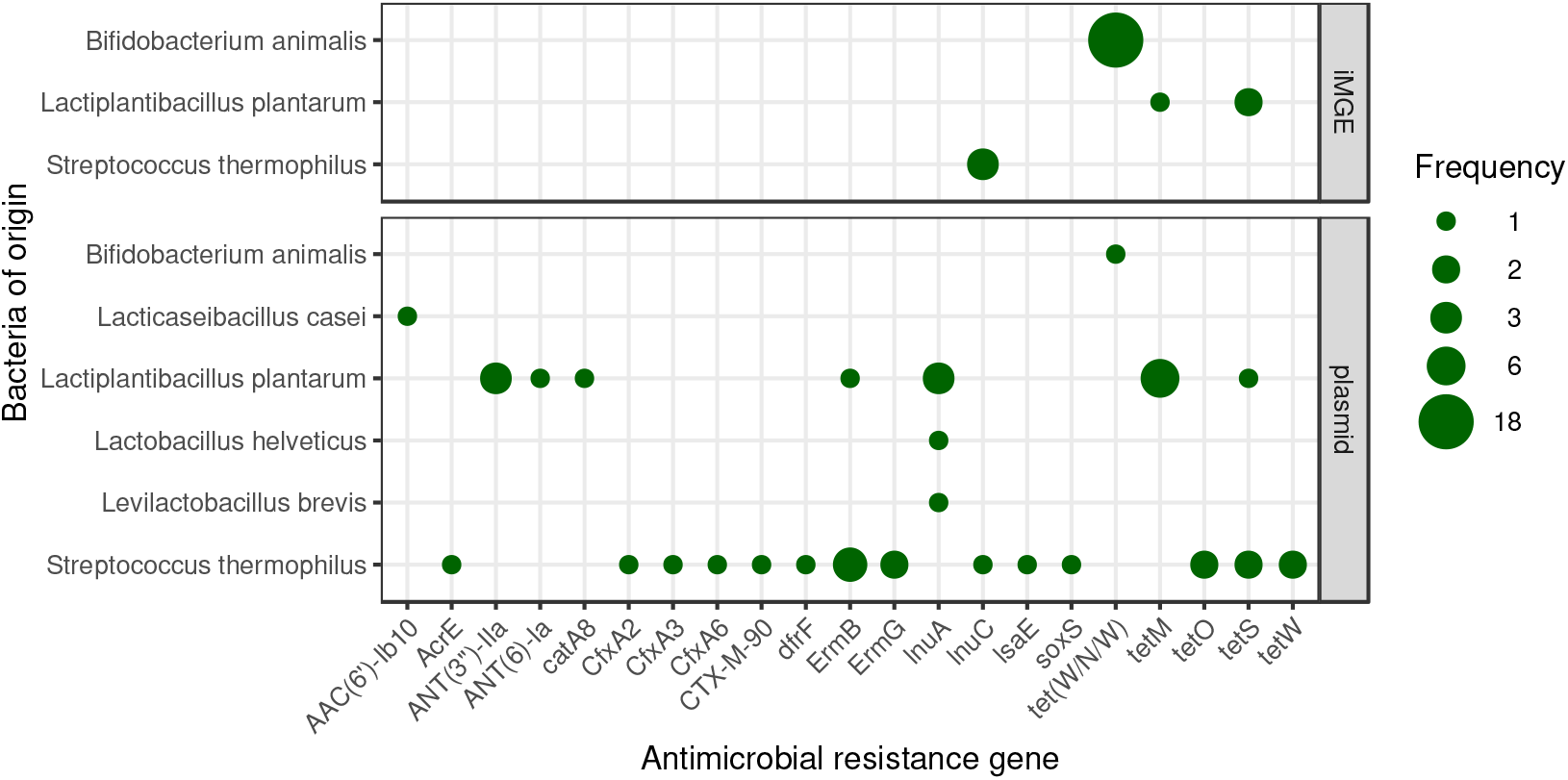
Mobile ARG frequency by bacteria of origin. The size of the dot indicates the occurrence frequency of the given gene flanked by iMGE or positioned in a plasmid. No ARGs were detected in phages.

#### iMGEs

Our results show that iMGE associated ARGs were detected in three species (Bifidobacterium animalis, Lactiplantibacillus plantarum and Streptococcus thermophilus). These iMGEs were physically close to the ARG in the chromosome, thus providing mobility potential. In 18 metagenomic samples we found tet(W/N/W) associated with iMGEs on contigs classified as Bifidobacterium animalis originated. In three further samples inuC is linked to an iMGE on Streptococcus thermophilus originated contigs. The ARGs tetM and tetS were also linked with iMGEs on Lactiplantibacillus plantarum originated contigs.

#### Plasmids

Using another technique, we predicted if contigs with ARGs were coming from plasmids. This prediction shows the number of times each gene occurred on plasmid sequences in the bacterial species. In Bifidobacterium animalis, we identified a plasmid associated contig with tet(W/N/W). We also identified a plasmid associated contig with AAC(6’)-lb10 classified as Lacticaseibacillus casei. In both the Lactobacillus helveticus and Levilactobacillus brevis samples, one contig of plasmid origin had the InuA gene. In the Lactiplantibacillus plantarum samples, the genes ANT(3”)-IIa, ANT(6)-Ia, catA8, ErmB, lnuA, tetM and tetS were detected in 3, 1, 1, 1, 3, 6 and 1 contigs of plasmids respectively. In the Streptococcus thermophilus samples a plasmid associated contig harboured each of the genes AcrE, CfxA2, CfxA3, CfxA6, CTX-M-90, dfrF, lnuC, lsaE and soxS. Furthermore, in the Streptococcus thermophilus samples, the genes ErmB, ErmG, tetO tetS and tetW were detected in 3, 2, 2, 2 and 2 contigs of plasmids respectively.

## Discussion

Our study confirms that numerous ARGs are present in kefir and yoghurt products and that many of them are mobile. Thus, yoghurt and kefir have the potential to contribute to AMR. Figure 1 shows the proportion of ARG positive samples by bacterial species. We found that Levilactobacillus brevis, Lactobacillus helveticus, Lactiplantibacillus plantarum and Lacticaseibacillus casei all had low proportions of ARG positive samples (<10%). Lactiplantibacillus plantarum is one of the most commonly used strains in probiotics so it is important to note that it had a low proportion of ARG positive samples indicating that it is a good choice for continued use in probiotics. This is in contrast with Bifidobacterium animalis which was found in fewer samples but had proportionally higher ARG positive samples with nearly all samples containing ARGs and thus our study suggests that it is not an optimum choice for probiotic use. In addition, no ARGs were found in Lactiplantibacillus argentoratensis and Lacticaseibacillus paracasei indicating that these are also good choices for probiotic cultures though there was only two samples of each so further studies would be needed to determine their suitability. In Table 1 we can see the ARGs found listed by species. It shows that although Bifidobacterium animalis had the highest proportion of ARGs (Figure 1), it had the lowest diversity of ARGs with only two distinct ARGs identified. Sixteen ARGs were identified in Lactiplantibacillus plantarum but it is important to bear in mind that it had by far the largest sample size which may skew results. Interestingly, the same five ARGs were identified in both Levilactobacillus brevis and Lactobacillus helveticus and these were the only ARGs found in both species. We identified a further four ARGs in Lacticaseibacillus casei. Furthermore, twenty-six distinct ARGs were identified in Streptococcus thermophilus making its ARG content the most diverse. In addition, in Figure 2 we can see that the number of distinct ARGs found within a given sample ranged from 1 to 20 with the highest number of ARGs being found in sample SRR11277066 from Streptococcus thermophilus.

In Figure 3 we identified the mobile ARG frequencies by bacterial species. The mobility potentials of the ARGs were predicted based on identifying iMGEs and plasmids as these may play a significant role in HGT. Where iMGEs are identified in the sequence context of an ARG, greater mobility can be assumed. The case is the same if the contig harbouring an ARG is plasmid originated. There were 18 occurrences of tet(W/N/W) where iMGEs were close to the ARG in the chromosome and one incidence where it was positioned in a plasmid. This was by far the most frequently identified mobile ARG but the only bacterial species it was found in was Bifidobacterium animalis. It is also important to note that despite its abundance, it was the only mobile ARG we could find in Bifidobacterium animalis. In contrast, in Streptococcus thermophilus twenty occurrences of plasmid originated ARGs were identified and three of ARGs being flanked by iMGEs, totalling twenty-three occurrences of mobile ARGs from fourteen distinct ARGs. Lactiplantibacillus plantarum had three incidences of ARGs being flanked by iMGEs and sixteen of plasmid originated ARGs, totalling nineteen incidences of mobile ARGs from nine distinct ARGs. Six of these mobile ARG occurrences were the TetM gene thus making it the ARG with the second highest mobility potential. Lacticaseibacillus casei, Levilactobacillus brevis and Lactobacillus helveticus all had one incidence of plasmid originated ARG with the same ARG identified in Levilactobacillus brevis and Lactobacillus helveticus, InuC.

In Table 1 we listed the ARGs found by species. The ARGs found in Bifidobacterium animalis (Bifidobacterium adolescentis rpoB mutants conferring resistance to rifampicin and tet(W/N/W)) have been previously identified as occurring in this species. To our knowledge, none of the four ARGs found in Lacticaseibacillus casei (AAC(6’)-Ib10, AAC(6’)-Ic, ANT(3”)-IIa and Escherichia coli EF-Tu mutants conferring resistance to Pulvomycin) have been previously identified as occurring in Lacticaseibacillus casei in the literature associated with this topic. Also, of the ARGs found in Lactiplantibacillus plantarum (AAC(6’)-Ii, ANT(3”)-IIa, ANT(6)-Ia, Lactobacillus reuteri cat-TC, catA8, eatAv, ErmB, lnuA, msrC, patA, patB, pmrA, poxtA, RlmA(II), tetM and tetS), none are identified in the literature as occurring in this species according to our knowledge. Furthermore, none of the ARGs found in Lactobacillus helveticus (lnuA, patA, patB, pmrA and RlmA(II)) nor in Levilactobacillus brevis (lnuA, patA, patB, pmrA and RlmA(II)) have been previously identified as occurring in these species as far as I have been able to ascertain. In addition, according to our knowledge this is the first time the ARGs found in Streptococcus thermophilus (AAC(6’)-Ie-APH(2”)-Ia, AAC(6’)-Ii, aadS, AcrE, APH(2”)-IVa, CblA-1, CfxA2, CfxA3, CfxA6, CTX-M-90, dfrF, eatAv, ErmB, ErmF, ErmG, FosA3, lnuC, lsaE, soxS, tet(40), Tet(X1), tet32, tetO, tetQ, tetS, tetW and tetX) have been identified in this species.

In Figure 2 we can see all samples where ARGs were detected. This shows that the most common ARGs were rpoB mutants conferring resistance to rifampicin and tet(W/N/W) detected in 36 and 20 samples respectively, all in Bifidobacterium animalis. Bifidobacterium adolescentis rpoB mutants conferring resistance to rifampicin is a rifamycin-resistant beta-subunit of RNA polymerase (rpoB). It has the resistance mechanisms of antibiotic target alteration and antibiotic target replacement. It is capable of conferring resistance to the Rifamycin drug class. In the literature this gene is published as being found in the following bacteria; Bifidobacterium animalis, Bifidobacterium breve, Bifidobacterium longum, Bifidobacterium thermophilum, Gardnerella vaginalis and Streptococcus pneumoniae. Similarly to what we found in the literature, we also identified it as existing in Bifidobacterium animalis. We identified tet(W/N/W) as having the highest mobility potential of all ARGs found. tet(W/N/W) is a mosaic tetracycline resistance gene and ribosomal protection protein. It has the resistance mechanism of antibiotic target protection and has an effect on tetracylines. In the literature this gene has been identified in a large variety of bacterial species. Our finding is similar to the previous findings published by other authors as we also found it in Bifidobacterium animalis.

The ARGs we identified may undermine several classes of antibiotics such as rifamycin, tetracycline, aminoglycoside, elfamycin, phenicol, fluoroquinolone, lincosamide, macrolide, oxazolidinone, pleuromutilin, streptogramin, carbapenem, cephalosporin, cephamycin, diaminopyrimidine, fosfomycin, glycylcycline, monobactam, penam, penem and triclosan. The ARGs we found have resistance mechanisms against some of the most important antibiotics, both in human and animal medicine. The term critically important antimicrobial (CIA) refers to antimicrobials which are last resorts in the treatment of human disease (DAFM, 2018). The WHO produces an updated list of currently used human antimicrobials grouped under three categories according to their importance; CIA, Highly important antimicrobial (HIA) and important antimicrobial (IA). CIAs are further subdivided into high priority CIA (CIA) and highest priority CIA (HPCIA). Most importantly are those listed as HPCIA which includes cephalosporins (3rd, 4th and 5th generation), glycopeptides, macrolides and ketolides, polymyxins and quinolones (WHO, 2018a). Out of the five HPCIA drug groups, we found ARGs which compromise the effectiveness of three (fluoroquinolones, cephalosporins and macrolides). We also found ARGs which have an effect on eight CIAs, five HIAs and one IA. The EMA also produced a list aimed at restricting the veterinary use of antimicrobials which are important for human medicine (EMA, 2019a). The antimicrobials are listed under the categories; Avoid, Restrict, Caution and Prudence. We found ARGs that threaten eight drug groups listed as avoid, one listed as restrict, six as caution and one as prudence. In addition, the World Organisation for Animal Health (OIE) has a list of critically important antimicrobial agents used in veterinary medicine. The OIE uses three categories; Veterinary Critically Important Antimicrobial Agents (VCIA), Veterinary Highly Important Antimicrobial Agents (VHIA) and Veterinary Important Antimicrobial Agents (VIA). The ARGs we found have an effect on five VCIAs, five VHIAs and one VIA. Thus, many of the most important antibiotics in human and animal medicine could be affected by the ARGs we detected in bacterial strains from kefir and yoghurt products.

The results of our study indicate that the use of bacteria such as Bifidobacterium animalis and Streptococcus thermophilus in probiotics should be reconsidered due to their high ARG content. Streptococcus thermophilus had the highest diversity of ARGs as well as the highest abundance and diversity of mobile ARGs. Bifidobacterium animalis had the lowest diversity of ARGs but the highest proportion of ARG positive samples with nearly all samples containing ARGs. It was also the only bacterial species to contain the ARG which we found to have the highest mobility potential, tet(W/N/W). Thus, our study suggests that these are not optimum choices for continued use in probiotic cultures. In contrast, our findings suggest that the use of other species such as Lactiplantibacillus plantarum may be better choices for continued use. Lactiplantibacillus plantarum is one of the most commonly used strains in probiotics so it is important to note that it had a low proportion of ARG positive samples. However, we did identify sixteen distinct ARGs in addition to nineteen occurrences of mobile ARGs in Lactiplantibacillus plantarum, though the large sample size may somewhat skew results. No ARGs were found in Lactiplantibacillus argentoratensis and Lacticaseibacillus paracasei indicating that these may be good choices for probiotic cultures, though there were only two samples of each so further studies with larger sample sizes would be needed to determine their suitability. The same ARGs and the same mobile ARG were identified in both Levilactobacillus brevis and Lactobacillus helveticus indicating they have similar effects on the resistome and mobilome. Levilactobacillus brevis, Lactobacillus helveticus and Lacticaseibacillus casei all had low proportions of ARG positive samples indicating they are potentially good options for use in probiotics. Further studies assessing the ARG diversity, frequency and mobility of bacterial strains used in fermented foods are needed to better assess the danger of the currently used strains and to explore potential alternative bacterial strains suitable for use. In this study, I curated the data and selected and downloaded the appropriate datasets from the SRA database; this could also be extended by further bioinformatic steps.

In conclusion, our study highlights the need for starting cultures of probiotics, such as yoghurt and kefir products, to be strictly monitored and bacteria of low ARG content selected for use. We found numerous and diverse ARGs in commonly used bacterial strains of kefir and yoghurt products. Thus, the results of our study support the findings of several other recent studies which have also identified ARGs in probiotics^5, 6, 33–35^. We also found that many of them have the potential to be mobile. Thus, fermented foods can act not only as a reservoir for ARGs but also as a medium for their exchange. During the fermentation process, the ARG content of yoghurt and kefir increases and, with the aid of plasmids and iMGEs such as we found, a potential hotspot for AMR development is created. Given the popularity of probiotics worldwide and the urgent threat of AMR, it is of utmost importance to fully investigate the risks associated with the consumption of probiotics. Considering the direct interactions humans have with animals, the interactions with the environment and the consumption of these animals and their produce, the implementation of a one health approach is needed. The prudent use of antimicrobials in human medicine and the strict monitoring of antimicrobial use in livestock with the aim of reduction is vital. In addition to medications, investigating potential environmental and animal sources of ARGs is essential to help tackle this threat for the sake of both veterinary and human medicine. Our study helps bring to light the fact that foods entering the body should be regarded as potential sources of ARGs, especially in light of the important classes of drugs these ARGs found are known to affect. Thus, going forward, there is a need for further studies with larger sample sizes of more commonly and lesser used probiotic bacterial strains.

## Supplementary materials

**Table 2.**
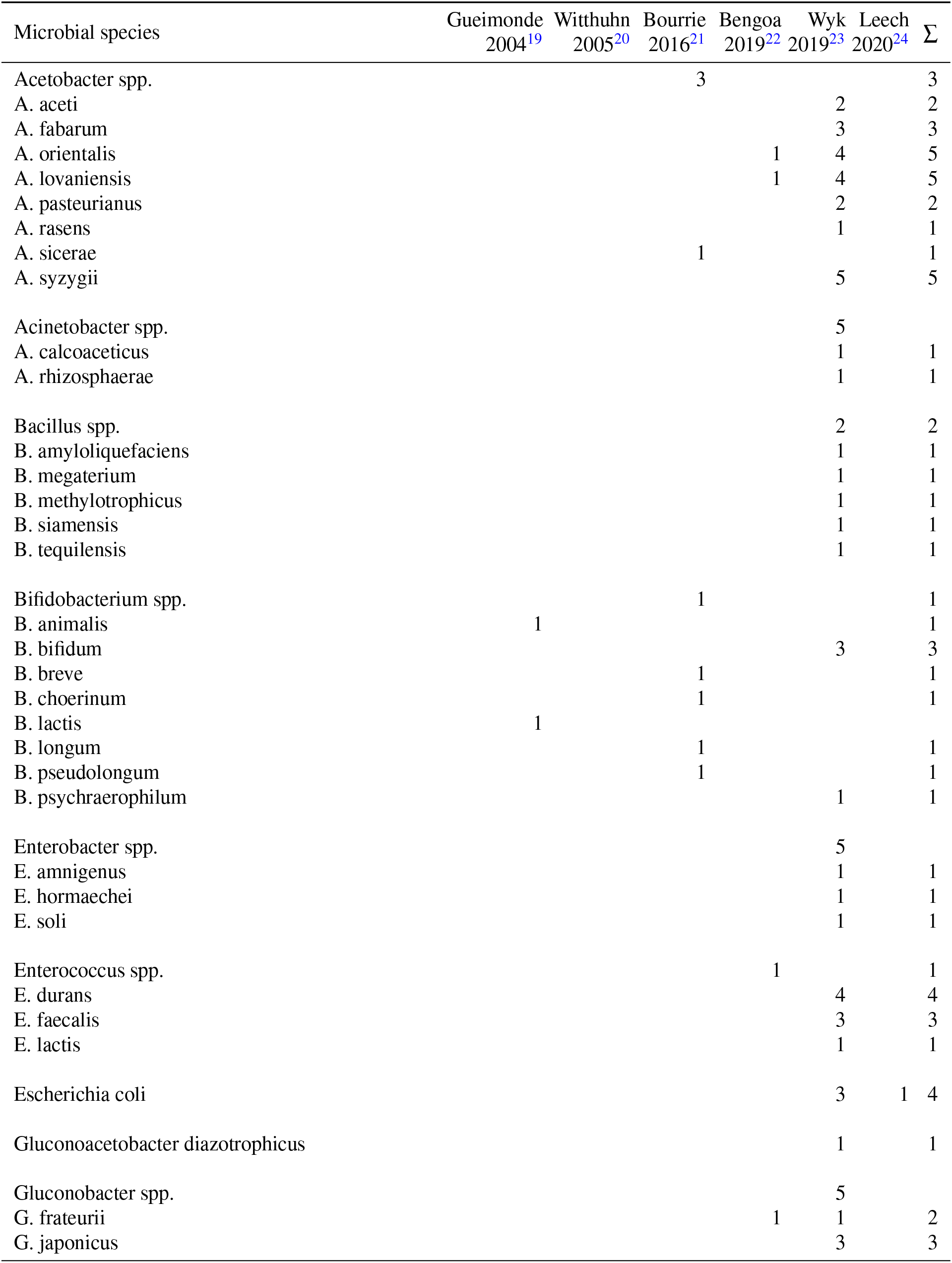
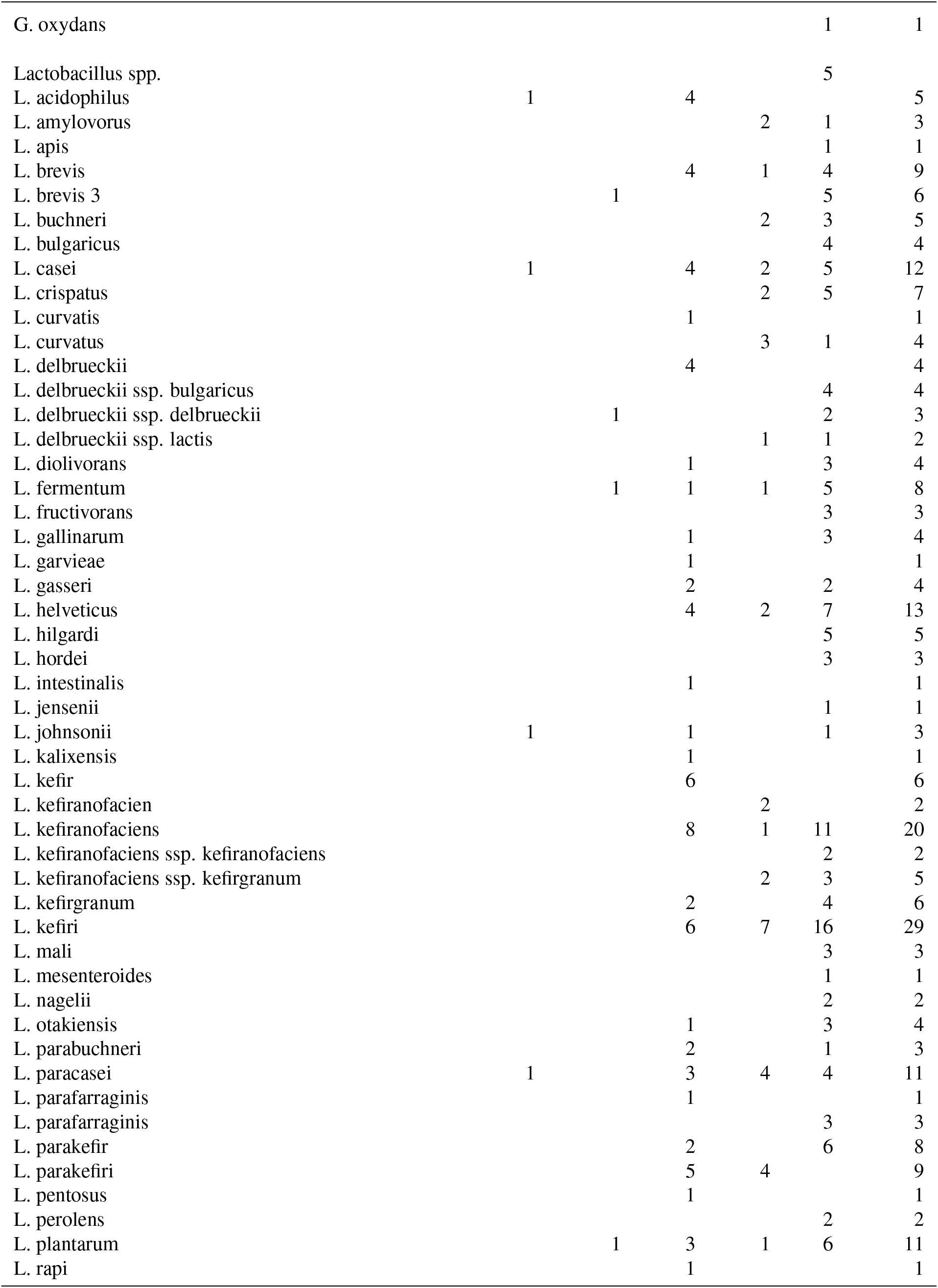
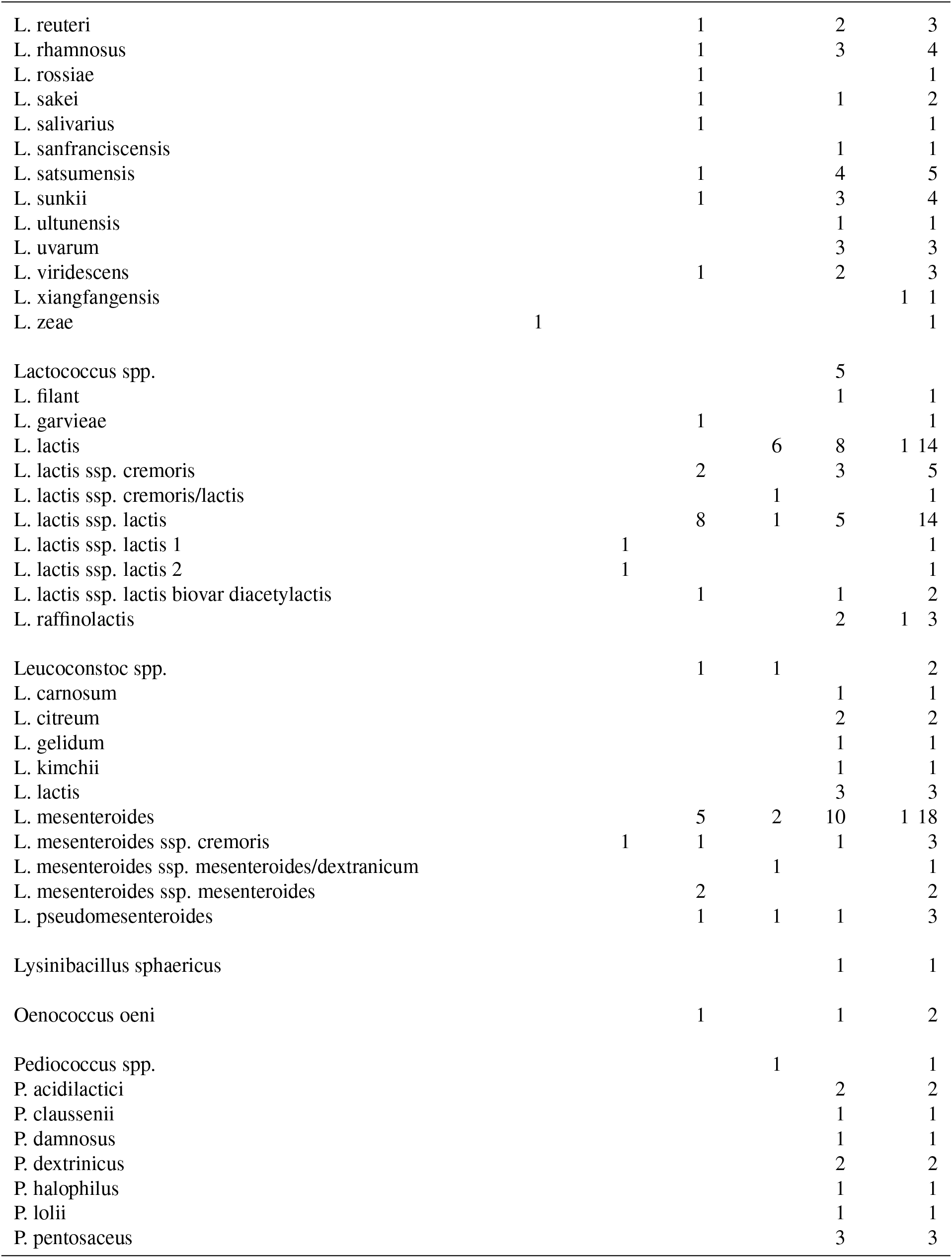
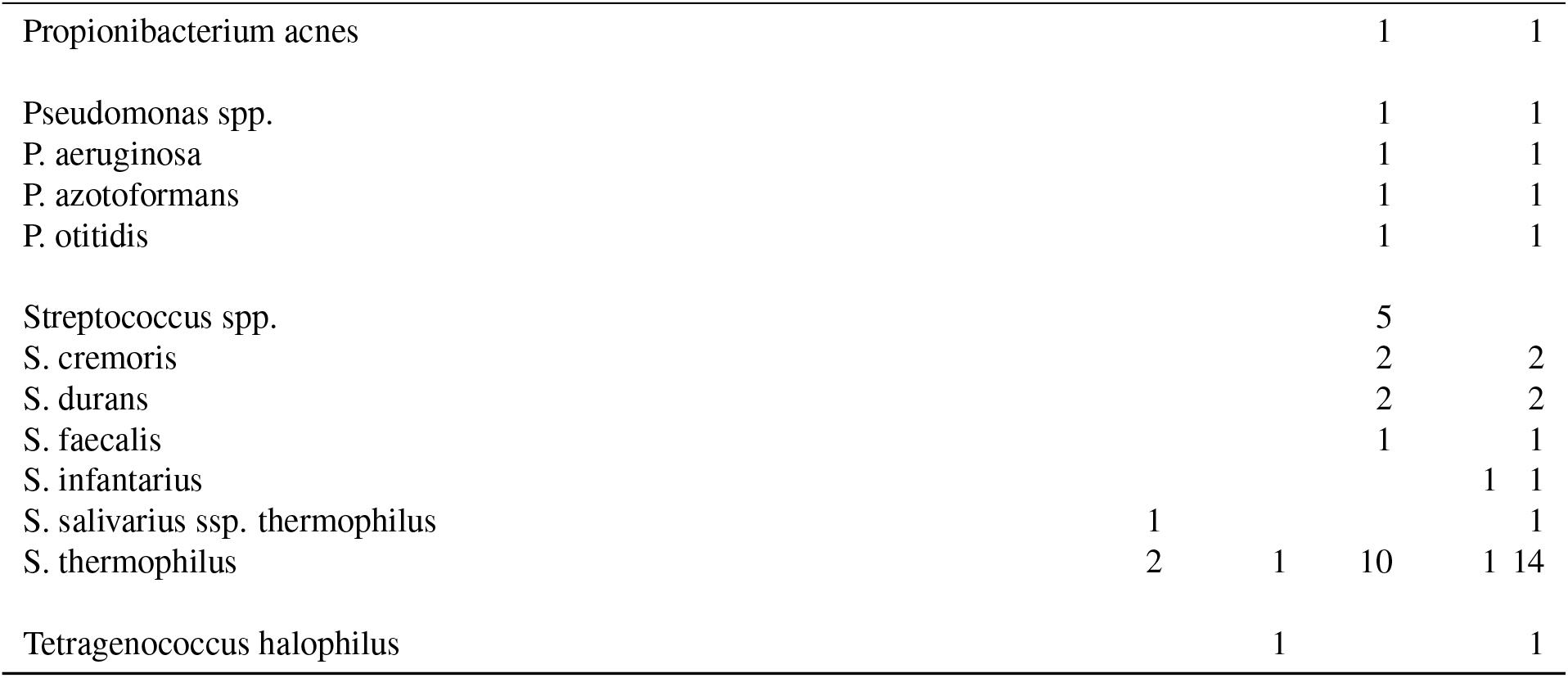
The number of studies that identified each bacterial species as occurring in kefir or yoghurt products. Duplicate sources were omitted.

### Selected NCBI SRA runs

The identifiers of the analysed samples (n=584) obtained from the NCBI SRA repository by species are the following:

*Bifidobacterium animalis*: ERR2221337, ERR2221385, ERR2397402, ERR3931553, ERR3931556, ERR3931557, ERR3931563, ERR3931572, ERR3931573, ERR3931581, ERR3931588, ERR3931756, ERR3931758, ERR4552599, ERR4552600, ERR4552601, ERR4552602, ERR4552603, ERR4552604, SRR11515245, SRR2124893, SRR5310856, SRR5310858, SRR5310869, SRR5310872, SRR5310875, SRR7030678, SRR7030680, SRR7030681, SRR7030682, SRR7030689, SRR7102022, SRR8060796, SRR8382541, SRR9274925, SRR9275544, SRR9275545.

*Lacticaseibacillus casei*: SRR3944187, SRR3944188, SRR5518762, SRR5518764, SRR5518765, SRR5518846, SRR5518847, SRR5518848, SRR5518849, SRR5518850, SRR5518851, SRR5518852, SRR5518853, SRR5518854, SRR5518855, SRR5518856, SRR5518857, SRR5518858, SRR6790308, SRR6790309, SRR6790310, SRR6790312, SRR6790313, SRR6790314, SRR6790315, SRR6790316, SRR6790317, SRR6790318, SRR6790319, SRR6790320, SRR6790321, SRR6790322, SRR6790323, SRR6790324, SRR6790325, SRR6790326, SRR6790327, SRR6790328, SRR6790329, SRR6790330, SRR6790331, SRR6790332, SRR6790333, SRR6790334, SRR6790335, SRR6790336, SRR6790337, SRR6790338, SRR6790339, SRR6790340, SRR6790341, SRR6790342, SRR6790343, SRR6790344, SRR6790345, SRR6790346, SRR6790347, SRR6790348, SRR6790349.

*Lacticaseibacillus paracasei*: SRR3944189, SRR3944190.

*Lactiplantibacillus argentoratensis*: ERR387522, SRR1151228.

*Lactiplantibacillus plantarum*: ERR1158396, ERR1158397, ERR1554589, ERR1554590, ERR1554591, ERR2221349, ERR2286790, ERR2286794, ERR2286891, ERR2286898, ERR2286899, ERR298627, ERR298635, ERR298703, ERR298714, ERR3151426, ERR3159182, ERR3162803, ERR3162986, ERR3283901, ERR3330792, ERR3330865, ERR3330866, ERR3330867, ERR3330868, ERR3330869, ERR3330870, ERR3330937, ERR386058, ERR386059, ERR3899072, ERR433488, ERR4593526, ERR4593527, ERR4593528, ERR4593530, ERR4593549, ERR4593550, ERR4833518, ERR485022, ERR485030, ERR485098, ERR485109, ERR570145, ERR570151, ERR570177, ERR570181, ERR570183, ERR570281, ERR570284, ERR570285, SRR10291920, SRR10357779, SRR10442245, SRR10605764, SRR10605765, SRR10605766, SRR10605767, SRR10605768, SRR10605769, SRR10605770, SRR10605771, SRR10605772, SRR10605773, SRR10605774, SRR10605775, SRR10605776, SRR10605777, SRR10605778, SRR10605779, SRR10605780, SRR10605781, SRR10605782, SRR10605783, SRR10605784, SRR10605785, SRR10605786, SRR10605787, SRR10605788, SRR10605789, SRR10605790, SRR10605791, SRR10605792, SRR10605793, SRR10605794, SRR10605795, SRR10605796, SRR10605797, SRR10605798, SRR10605799, SRR10605800, SRR10605801, SRR10605802, SRR10605803, SRR10605804, SRR10605805, SRR10605806, SRR10605807, SRR10605808, SRR10605809, SRR10605810, SRR10605811, SRR10605812, SRR10605813, SRR10605814, SRR10605815, SRR10605816, SRR10605817, SRR10605818, SRR10605819, SRR10605820, SRR10605821, SRR10605822, SRR10605823, SRR10605824, SRR10605825, SRR10605826, SRR10605827, SRR10605828, SRR10605829, SRR10605830, SRR10605831, SRR10605832, SRR10605833, SRR10605834, SRR10605835, SRR10605836, SRR10605837, SRR10605838, SRR10605839, SRR10605840, SRR10605841, SRR10605842, SRR10605843, SRR10605844, SRR10605845, SRR10605846, SRR10605847, SRR10605848, SRR10605849, SRR10605850, SRR10605851, SRR10605852, SRR10605853, SRR10605854, SRR10605855, SRR10605856, SRR10605857, SRR10605858, SRR10605859, SRR10605860, SRR10605861, SRR10605862, SRR10605863, SRR10605864, SRR10605865, SRR10605866, SRR10605867, SRR10605868, SRR10605869, SRR10605870, SRR10605871, SRR10605872, SRR10671639, SRR10671640, SRR10671641, SRR10671642, SRR10902843, SRR1151193, SRR11910127, SRR11910128, SRR11910136, SRR11910143, SRR11910158, SRR11910162, SRR11910168, SRR11910178, SRR11910187, SRR11910196, SRR11910200, SRR11910204, SRR11910205, SRR11910217, SRR11910220, SRR11910248, SRR11910255, SRR11910257, SRR11910269, SRR11910275, SRR11910286, SRR11910296, SRR11910298, SRR11910303, SRR11910305, SRR11910311, SRR11910337, SRR11910344, SRR11910354, SRR11910358, SRR11910365, SRR11910395, SRR11910414, SRR11910423, SRR11910429, SRR11910437, SRR12145645, SRR12559458, SRR12559459, SRR12559460, SRR12559461, SRR12559462, SRR12559464, SRR12559465, SRR12559467, SRR12559468, SRR12559469, SRR12559470, SRR12559471, SRR12559472, SRR12559473, SRR12559474, SRR12559476, SRR12559477, SRR12559478, SRR12559479, SRR12559480, SRR12559481, SRR12559482, SRR12559483, SRR12559484, SRR12559485, SRR12559487, SRR12559488, SRR12559489, SRR12559490, SRR12559491, SRR12559707, SRR12559708, SRR12559710, SRR12559711, SRR12559712, SRR12559713, SRR12559714, SRR12559715, SRR12559716, SRR12559717, SRR12559718, SRR12559719, SRR12559721, SRR12559722, SRR12559723, SRR12559724, SRR12559725, SRR12559726, SRR12559727, SRR12559728, SRR12559729, SRR12559730, SRR12559732, SRR12559733, SRR12559734, SRR12559735, SRR12559736, SRR12559737, SRR12559738, SRR12559739, SRR12559740, SRR12559741, SRR12559743, SRR12559744, SRR12559745, SRR12559746, SRR12559747, SRR12559748, SRR12559749, SRR12559750, SRR12559751, SRR12559752, SRR12559755, SRR12559756, SRR12559757, SRR12559758, SRR12559759, SRR12559858, SRR12559859, SRR12559860, SRR12559861, SRR12559862, SRR12559864, SRR12559865, SRR12559867, SRR12559868, SRR12559869, SRR12559870, SRR12559871, SRR12559872, SRR12559873, SRR12559875, SRR12559876, SRR12559877, SRR12559878, SRR12559879, SRR12559880, SRR12559881, SRR12559882, SRR12559883, SRR12559884, SRR12559886, SRR12559887, SRR12559888, SRR12559889, SRR12559890, SRR12560026, SRR12560027, SRR12560028, SRR12560029, SRR12560030, SRR12560032, SRR12560033, SRR12560034, SRR12875629, SRR13060588, SRR13213118, SRR13442537, SRR13442538, SRR13442539, SRR1552084, SRR1552590, SRR1552611, SRR1552612, SRR1552613, SRR1552614, SRR1552615, SRR1552616, SRR2142235, SRR4124949, SRR4124950, SRR4124954, SRR4124955, SRR5518758, SRR5518859, SRR5518863, SRR5518864, SRR5518865, SRR5518866, SRR5518867, SRR5518868, SRR5518869, SRR5518870, SRR5518871, SRR5518872, SRR5518873, SRR5518874, SRR5518875, SRR5724505, SRR5914586, SRR7550999, SRR7551002, SRR7551003, SRR7551004, SRR7551010, SRR8182735, SRR8252890, SRR8252891, SRR8382543, SRR8693953, SRR9107669, SRR9861755.

*Lactobacillus helveticus*: ERR204044, ERR298639, ERR298656, ERR298711, ERR3283916, ERR387534, ERR485034, ERR485051, ERR485106, SRR10332348, SRR1151128, SRR11910141, SRR11910150, SRR11910390, SRR12560068, SRR12560069, SRR4450492, SRR5724508, SRR9866100, SRR9866101, SRR9866102, SRR9866103, SRR9866104, SRR9866105, SRR9866106, SRR9866107, SRR9866108, SRR9866109, SRR9866110, SRR9866111, SRR9866112, SRR9866113, SRR9866114, SRR9866115, SRR9866116, SRR9866117, SRR9866118, SRR9866119, SRR9866120, SRR9866121, SRR9866122, SRR9866123, SRR9866124, SRR9866125, SRR9866126, SRR9866127, SRR9866128, SRR9866129, SRR9866130, SRR9866131, SRR9866132, SRR9866133, SRR9866134, SRR9866135, SRR9866136, SRR9866137, SRR9866138, SRR9866139, SRR9866140, SRR9866141.

*Levilactobacillus brevis*: ERR2305654, ERR3283759, ERR386054, SRR1151178, SRR11910179, SRR11910198, SRR11910213, SRR11910306, SRR11910310, SRR11910336, SRR11910381, SRR11910417, SRR11910436, SRR11910439, SRR12559992, SRR12559993, SRR12560016, SRR4450488, SRR5724506, SRR5724507, SRR5724509, SRR6918185, SRR6918186, SRR6918187, SRR6918188, SRR6918189, SRR6918190, SRR7551000, SRR7551001.

*Streptococcus thermophilus*: ERR3330766, SRR11276977, SRR11277033, SRR11277066, SRR11910208, SRR11910216, SRR11910219, SRR11910241, SRR11910242, SRR11910258, SRR11910321, SRR11910326, SRR11910350, SRR11910360, SRR11910376, SRR11910392, SRR11910418, SRR12037890, SRR5310871, SRR5310876, SRR5310877, SRR6319282, SRR6319283, SRR6319284, SRR6319285, SRR6319286, SRR7850709.

